# Conserved N-terminal Regulation of the ACA8 Calcium Pump with Two Calmodulin Binding Sites

**DOI:** 10.1101/2023.12.07.570580

**Authors:** Sigrid Thirup Larsen, Josephine Karlsen Dannersø, Christine Juul Fælled Nielsen, Lisbeth Rosager Poulsen, Michael Palmgren, Poul Nissen

## Abstract

The autoinhibited plasma membrane calcium ATPase, ACA8 from *A. thaliana* has an N-terminal autoinhibitory domain. Calcium-bound calmodulin binding at two sites located at residues 42-62 and 74-96 relieves autoinhibition of ACA8 activity.

We investigated N-terminally truncated ACA8 constructs (WT, Δ20, Δ30, Δ35, Δ37, Δ40, Δ74 and Δ100) to explore the role of conserved motifs in the N-terminal segment preceding the calmodulin binding sites. Furthermore, we purified WT, Δ20- and Δ100-ACA8, tested activity *in vitro* and performed structural studies of purified Δ20-ACA8 stabilized in its native form to explore the mechanism of autoinhibition.

Through activity studies and a yeast complementation assay, we show that an N-terminal segment between residues 20 and 35, upstream of the calmodulin binding sites, is important for autoinhibition and the activation by calmodulin, and that a conserved Phe32 is essential for autoinhibition. Cryo-EM structure determination at 3.3 Å resolution of a beryllium fluoride inhibited form shows no autoinhibition, but a low-resolution structure for an E1 state indicates autoinhibitory domain binding consistent with the mutational studies and AlphaFold predicted structures.

## Introduction

By maintaining a very low concentration of intracellular calcium, eukaryotic cells establish an approximately 20,000-fold gradient between the exterior and interior. Control of the intracellular calcium level is highly important, and correct responses to calcium signals affect all aspects of cell life and death. Uncontrolled changes or collapse of the gradient to the outside calcium level are incompatible with life and elevated calcium levels lead to programmed cell death [1–3]. In plants, calcium signalling controls responses to abiotic cues such as light, temperature and mechanical stress, and biotic cues such as cellular signals or the activity of other living organisms [4–7]. Similarly, calcium signalling is associated with nearly all signalling and response pathways in mammals, including e.g. neuronal development and signalling, memory consolidation, circadian rhythms, hormone secretion, muscle contraction and fertility [1, 8, 9].

Calcium ATPases belonging to the P2B subgroup of the P-type ATPase family [10] play a vital role in maintaining a steep Ca^2+^ gradient with a tightly controlled level of intracellular calcium. They encompass a range of typically calmodulin activated Ca^2+^-ATPase including plasma membrane Ca^2+^-ATPases (PMCAs) in animals [11, 12], which are also alpha-synuclein activated [13], and autoinhibited Ca^2+^-ATPase (ACAs) in plants [14–16].

The mammalian PMCAs transport Ca^2+^ from the intracellular space to the confined and controlled extracellular environment and maintain the overall calcium homeostasis [12]. PMCAs represent the high-affinity Ca^2+^ system that extrudes smaller, but defining amounts of Ca^2+^ from the cell after a Ca^2+^ signalling event [11, 12], whereas larger volume Ca^2+^ extrusion is typically performed by the sodium-calcium exchanger (NCX) or removed by reuptake to internal stores by sarco/endoplasmic reticulum Ca^2+^-ATPase (SERCA). Potent PMCA activity may also elicit local control by active calcium extrusion and calcium depression [17, 18].

The ACAs of plants do not localise exclusively to the plasma membrane. In *Arabidopsis thaliana* there are ten different ACAs with six of them known to localize to the plasma membrane, including ACA8 [19–23], while others are localized to endomembrane systems such as the endoplasmic reticulum and vacuoles [24–26]. The ACAs are subject to a complex and dynamic network of posttranslational regulation and are essential for the plants’ response to the environment [27].

A hallmark of both PMCAs and ACAs is an autoinhibitory domain important for regulation of pump activity. Interestingly, the autoinhibitory domain is situated C-terminally for mammalian PMCA and N-terminally for ACAs in plants [28–31]. However, the autoinhibition of both ACA8 and mammalian PMCA is independent of the position of the autoinhibitory domain [28, 32], and the respective mechanisms appear to be related even though the evolution of their autoinhibitory domains is independent of each other [33].

In the resting state, the autoinhibitory domain interacts with the intracellular domains thereby preventing ATPase activity and transport (or backflow) of Ca^2+^ across the membrane [29, 31]. Upon an increase in intracellular calcium, Ca^2+^-loaded calmodulin (here just denoted CaM) will bind to the autoinhibitory domain and relieve autoinhibition [11, 34].

In 2004 Luoni et al. showed that not only a canonical CaM binding site (residue 41-63) of the autoinhibitory domain of *A. thaliana* ACA8 was important for autoinhibition; the full-length N-terminal segment (1-116) inhibited activity to a higher degree than the CaM binding site alone [35]. An explanation was ascribed later to a second CaM binding site (at residues 74-96) identified by Tidow et al., and a two-step activation mechanism was revealed on the basis of structural and biochemical studies of the isolated autoinhibitory domain in complex with Ca^2+^-CaM combined with ACA8 complementation studies in a yeast Ca^2+^-ATPase knock-out strain [36].

From truncation studies presented here, we now find that also a conserved stretch, located N terminally to the two CaM binding sites in ACA8, is important for autoinhibition. Based on sequence alignments, the N-terminal truncation constructs Δ20-, Δ30-, Δ35-, Δ37- and Δ40-ACA8 (from now on referred to by their truncation site only), were created and expressed in yeast, and the protein-containing membranes were tested for their Ca^2+^ and CaM dependent activity. The constructs were compared to WT ACA8 and to the Δ74 and Δ100 constructs that are known to lack autoinhibition [36]. The impact of N-terminal truncations was also studied through an *in vivo* assay, and on purified WT, Δ20 and Δ100 through their Ca^2+^ and CaM dependent ATPase activity. The truncation studies further motivated investigations, both *in vitro* and *in vivo,* of an F32A mutant for WT and Δ20. Finally, cryo-EM studies were performed of the autoinhibited Δ20 in two functional states. Combined with an AlphaFold predicted structure, it leads us to a detailed model of the autoinhibitory mechanism for ACA8.

## Results

A series of truncation constructs of ACA8 (Figure 1) were created and expressed in the calcium pump deficient yeast strain K616. Δ40 is the shortest construct preserving both CaM binding sites (CaMBS) [36]. The Δ20 construct preserves a stretch of conserved residues of the N-terminus, whereas the Δ30, Δ35 and Δ37 constructs contain progressively fewer of these (Figure 1A) [19]. Residues 1 to 19 are highly variable, generally of charged and polar character and predicted to be disordered (Figure S1).

**Figure 1:**
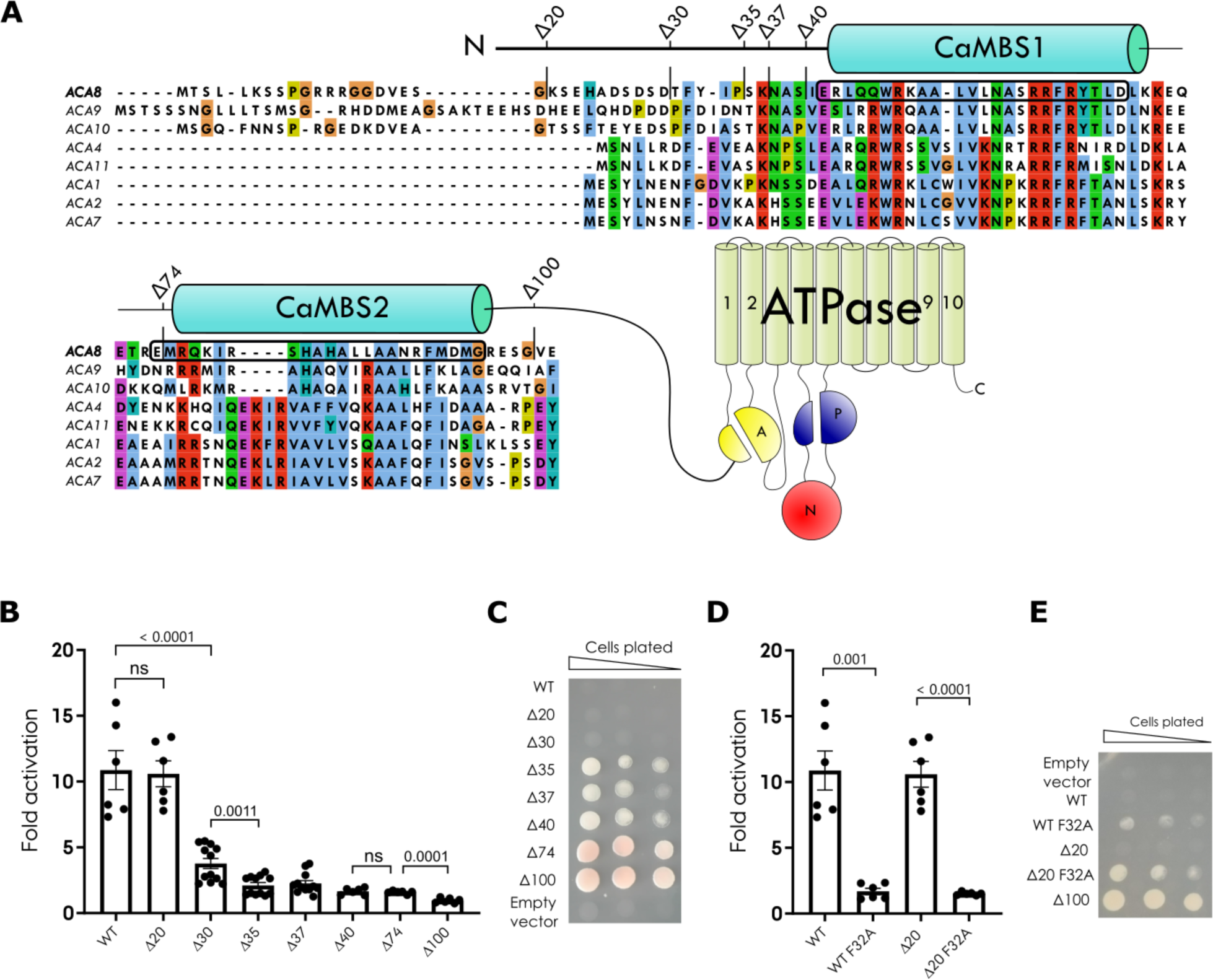
Sequence alignment, truncation constructs and activity of membrane bound ACA8 constructs. **A)** Sequence alignment of ACA8 from *A. thaliana* and 7 other ACA’s from *A. thaliana.* The protein sequence is coloured according to CLUSTAL colouring scheme. The start position of the different N-terminal truncation constructs is indicated for the sequence of ACA8 as well as the position of the CaM binding sites (CaMBS). The sequence alignment is visualised using Jalview [38]. **B)** CaM stimulated activity observed for the different membrane bound ACA8 constructs. The fold activation represents the increase in activity observed in the presence of CaM. Data presented as mean ± SEM. n = 6 for WT, Δ20, Δ40, Δ74, and Δ100 and n = 12 for Δ30 Δ35 and Δ37. The statistical analysis is conducted as an unpaired, two-tailed students t-test. Activity measurements were performed on two independent protein expressions in at least triplicates. **C)** Complementation assay, only cells expressing an active form of ACA8 can grow under Ca^2+^-deprived conditions. K616 yeast strain is devoid of calcium pumps and transformed with WT, Δ20, Δ30, Δ35, Δ37, Δ40, Δ74, Δ100 constructs of ACA8 (the latter as a positive control) and empty vector (negative control), respectively. The transformed cells were grown on medium with 2% galactose and 10mM EGTA (SGA EGTA). ACA8 construct expression is controlled by a galactose-inducible promoter, GAL1. Control plates (SGA CaCl_2_, SDA EGTA and SDA CaCl_2_) are shown in the supplementary material (Figure S4). **D)** CaM stimulated activity observed for the F32A ACA8 mutants. The fold activation represents the increase in activity observed in the presence of CaM. Data presented as mean ± SEM, n = 6 for all constructs. The statistical analysis is conducted as an unpaired two-tailed students t-test. Activity measurements were performed on two independent protein expressions in triplicates. **E)** Complementation assay, plated on SGA EGTA. K616 yeast strain transformed with WT, Δ20, WT F32A, Δ20 F32A, Δ100 and empty vector respectively. Control plates (SGA CaCl_2_, SDA EGTA and SDA CaCl_2_) are shown in the supplementary material (Figure S5).

The expression levels of all constructs were checked by SDS-PAGE analysis (Figure S2), and the CaM activation of the membrane-bound protein was determined.

Comparison of the CaM stimulated ATPase activities reveals that the conserved N-terminal segment preceding the first CaM binding site (CaMBS1) is important for autoinhibition (Figure 1B). Removing more than the first 20 residues impacts CaM activation of the ATPase activity. Compared to WT and Δ20, the constructs Δ30, Δ35, Δ37 and Δ40 suffer a partial loss of autoinhibition and show only a low activation by CaM, comparable to the activation by CaM observed for Δ74 lacking CaMBS1.

For Δ30, the CaM activation is reduced almost 3-fold compared to WT (3.8 vs. 10.9-fold activation, respectively), but increased compared to Δ35 and Δ37. Although there is only a small difference between the fold activation of Δ30 and Δ35 (3.8 vs 2.1 fold-activation) (Figure 1B and Table 1) this indicates that the residues from 30 to 35 could have an impact on autoinhibition and CaM stimulation.

**Table 1:** Fold activation upon CaM stimulation of membrane bound ACA8. Mean value ± SEM for the fold activation of the different truncation constructs and F32A

| *Construct* | *Fold activation* |
| --- | --- |
| WT | 10.9 ± 1.5 |
| Δ20 | 10.6 ± 1.0 |
| Δ30 | 3.8 ± 0.4 |
| Δ35 | 2.1 ± 0.2 |
| Δ37 | 2.3 ± 0.2 |
| Δ40 | 1.7 ± 0.1 |
| Δ74 | 1.6 ± 0.05 |
| Δ100 | 1.0 ± 0.08 |
| WT F32A | 1.7 ± 0.2 |
| Δ20 F32A | 1.5 ± 0.05 |

*In vivo* investigations was performed through a yeast complementation assay that relies on an active Ca^2+^-ATPase to fill internal stores under Ca^2+^-deprived conditions [20, 36, 37]. Cells expressing fully deregulated (active) Δ100 grow on Ca^2+^ depleted media whereas cells expressing the autoinhibited WT-ACA8 cannot.

Figure 1C shows that Δ100 and Δ74 allow for cell growth of K616 as has also previously been described [36]. Oppositely, WT, Δ20 and Δ30 do not complement the K616 strain and therefore must maintain an effectively autoinhibited state of the pump.

The constructs Δ35, Δ37 and Δ40 complement the cells at an intermediate level. Hence, autoinhibition of these constructs is compromised, but not entirely absent, and the partial loss of autoinhibition observed in the activity studies (Figure 1B and Figure S3) for Δ35, Δ37 and Δ40 is apparently sufficient to allow for partial complementation. Where the *in vitro* activity studies of Δ30 and Δ35 were similar, the *in vivo* assay clearly distinguishes these two constructs and shows the importance of residues 30-35 for autoinhibition (Figure 1C).

Figure 1A shows that the 30-35 segment is conserved, including the fully conserved F32. Two constructs with a F32A mutation were created, namely full-length F32A and Δ20-F32A. The activity assays clearly show that the F32A mutants lacks autoinhibition as the fold activation by CaM is now only around 1.5 (Figure 1D and Table 1). The impact on autoinhibition was also tested *in vivo* by the complementation assay, and again absence of autoinhibition is observed (Figure 1E).

WT and Δ20, that are autoinhibited in the membranes, and Δ100, which is not, were purified and tested to check if autoinhibition also holds for detergent solubilised ACA8. The WT, Δ20 and Δ100 constructs were purified by Ni^2+^ affinity chromatography and size exclusion chromatography in the detergent LMNG and yielded pure protein (Figure S6, Figure S7, and Figure S8). The purified proteins were tested for both calcium and calcium/CaM dependent activity.

Figure 2A show the specific activity of the purified WT, Δ20 and Δ100 with and without CaM. Purified WT and Δ20 are autoinhibited. WT has a low basal activity of 0.242 ± 0.0446 nmol Pi/min/µg and CaM activated to 2.251 ± 0.150 nmol Pi/min/µg (Table S1) corresponding to a 10-fold activation. Δ20 has an even lower basal activity (0.063 (± 0.0024) nmol Pi/min/µg), and CaM stimulated activity is now 19-fold increased (reaching 1.2 ± 0.072 nmol Pi/min/µg) (Table S1). The basal activity of Δ100 is high (1.10 ± 0.104 nmol Pi/min/µg) and similar to CaM stimulated activity, as expected (1.16 ± 0.125 nmol Pi/min/µg) (Table S1). In other words, Δ100 is active in the absence of CaM and to the same degree as CaM activated Δ20. The activity of purified Δ20 was also tested in the presence of acidic phospholipids, which are known to relieve autoinhibition [39]. With acidic phospholipids present, Δ20 loses autoinhibition, and it has a high basal activity (1.55 ± 0.124 nmol Pi/min/µg) similar to the activity observed for Δ100. The added response to CaM is now only 1.8-fold reaching 2.76 ± 0.330 nmol Pi/min/µg (Figure 2A and Table S1).

**Figure 2:**
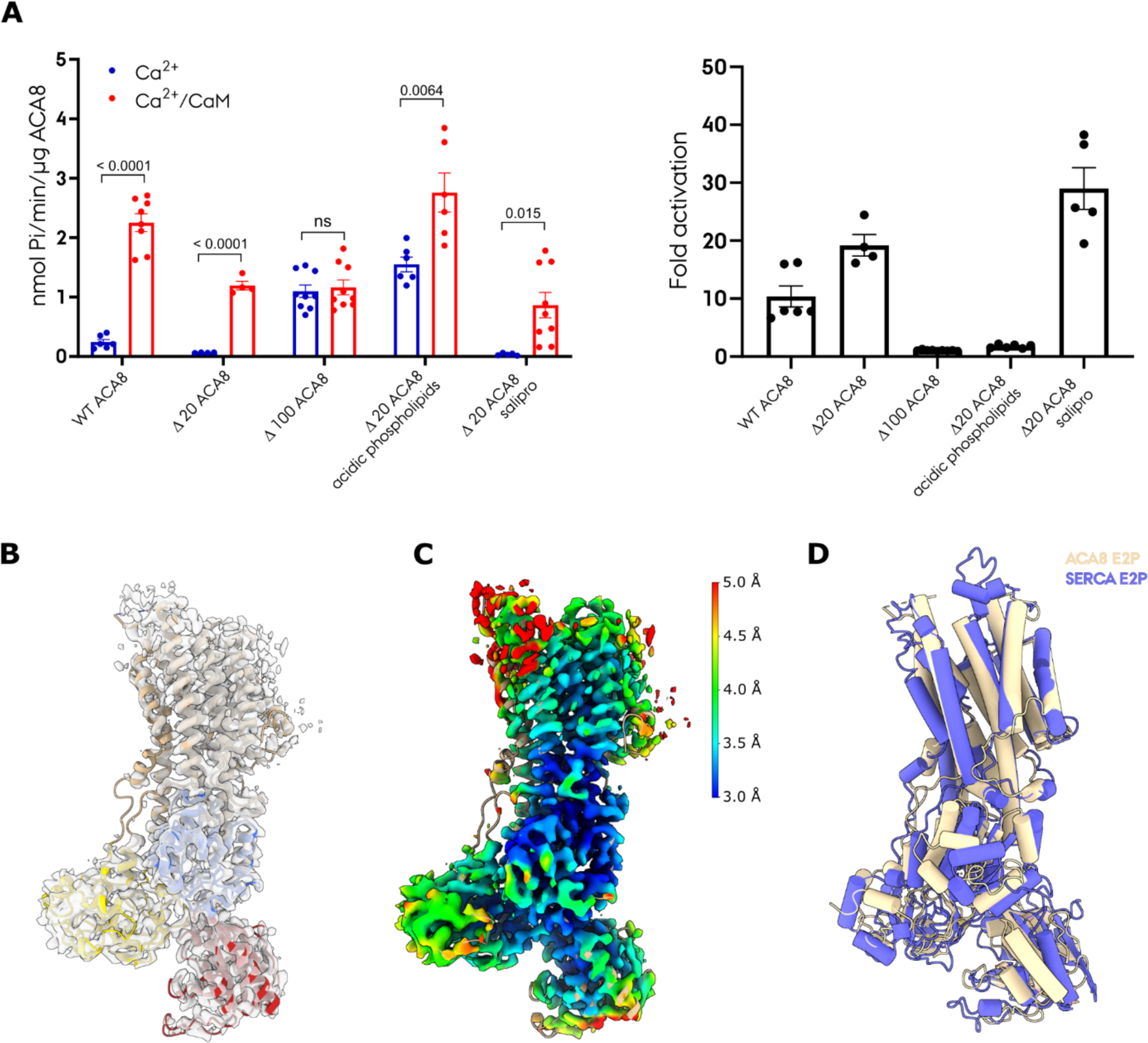
Activity and structure of purified ACA8. **A)** Activity measurements for purified ACA8. *Left:* Ca^2+^ dependent activity with and without CaM. Specific ATPase activity of purified WT, Δ20, Δ100 and of Δ20 with acidic phospholipids present and salipro reconstituted Δ20. WT n = 6 (Ca^2+^) and n = 8 (Ca^2+^/CaM), Δ20 n = 4, Δ100 n = 9, Δ20 acidic phospholipids n = 6, Δ20 salipro n = 5 (Ca^2+^) and n = 9 (Ca^2+/^CaM). Data presented as mean ± SEM. The statistical analysis is conducted as an unpaired two-tailed students t-test. *Right:* Fold activation of WT, Δ20, Δ100 and Δ20 activity with acidic phospholipids present and salipro reconstituted Δ20. Activity measurements are from at least two independent protein purification and expression cultures, except Δ100 which is from one expression culture. **B)** The 3.3Å (FSC gold standard = 0.143) density from cryo-EM in gray with the structure of ACA8 modelled. The transmembrane domain is wheat colored, the A domain is yellow, P domain is blue and N domain is red. Contour level 0.236. **C)** The density is colored by local resolution, from 3 Å (blue) to 5 Å (red). Contour level 0.236. **D)** The structure of ACA8, wheat colored, compared to E2P BeF structure of SERCA (PDB: **3B9B**) [40] in blue. Alignment on transmembrane helix 7 (TM7) to TM10.

Purified Δ20-ACA8 was reconstituted into saposin lipoprotein stabilized nanodiscs (salipro nanodiscs) [41, 42]. The ACA8 containing nanodiscs were separated from empty particles by size exclusion chromatography with a good separation and ratio of reconstituted protein and empty particles (Figure S9). The salipro reconstituted ACA8 also retained autoinhibition and CaM dependent activity (Figure 2A). ATPase activity is barely measurable in the absence of CaM, whereas the protein has an approx. 30-fold higher activity with CaM (Figure 2A and Table S1), reaching a comparable Ca^2+^ and CaM dependent activity (0.86 nmol Pi/min/µg) to what is observed for the Δ20 in LMNG (1.2 nmol Pi/min/µg) (Figure 2A and Table S1).

Using cryo-EM the Δ20-ACA8-salipro structure was determined in the presence of inhibitors mimicking specific functional states. BeFx was used to stabilize ACA8 in the E2P-like state, which stabilizes the autoinhibited state of related P4-ATPase lipid flippases of the P-type ATPase family [43].

A cryo-EM structure of BeFx-stabilized ACA8 was determined at 3.3 Å resolution overall. A model could be assigned for position 116 and onwards; but no density corresponding to the autoinhibitory domain could be observed (Figure 2B and C). As expected, the structure resembles an E2P state with the transmembrane domain (TM domain) as for the E2P state of SERCA [40] (Figure 2D), although with a slight vertical shift in the cytoplasmic domains (Figure S10). Unlike for P4-ATPases there is no indication of autoinhibition being associated with this state.

We then turned to AlphaFold for indications of how the autoinhibitory domain might interact with the core structure of ACA8.

AlphaFold predicts a structure for ACA8 with high confidence.

Figure 3A shows a cartoon representation of the predicted structure colored by confidence level from red/yellow (low confidence) to blue (high confidence). The first 24 residues are not confidently predicted with a score below 70, but from then on, including the autoinhibitory domain, the structure is confidently predicted with most residues having a pLDDT score of 70-100 (Figure S11). Importantly, the predicted structure shows an interaction of the autoinhibitory domain with the core of the pump (Figure 3A). The overall conformation of the predicted structure represents an E1 state, with an open access to the Ca^2+^ binding site as a clear characteristic. Comparing the core of the AlphaFold predicted structure with the PMCA1 structure in the E1 state [44], the TM domains superimpose well (Figure 3B).

**Figure 3:**
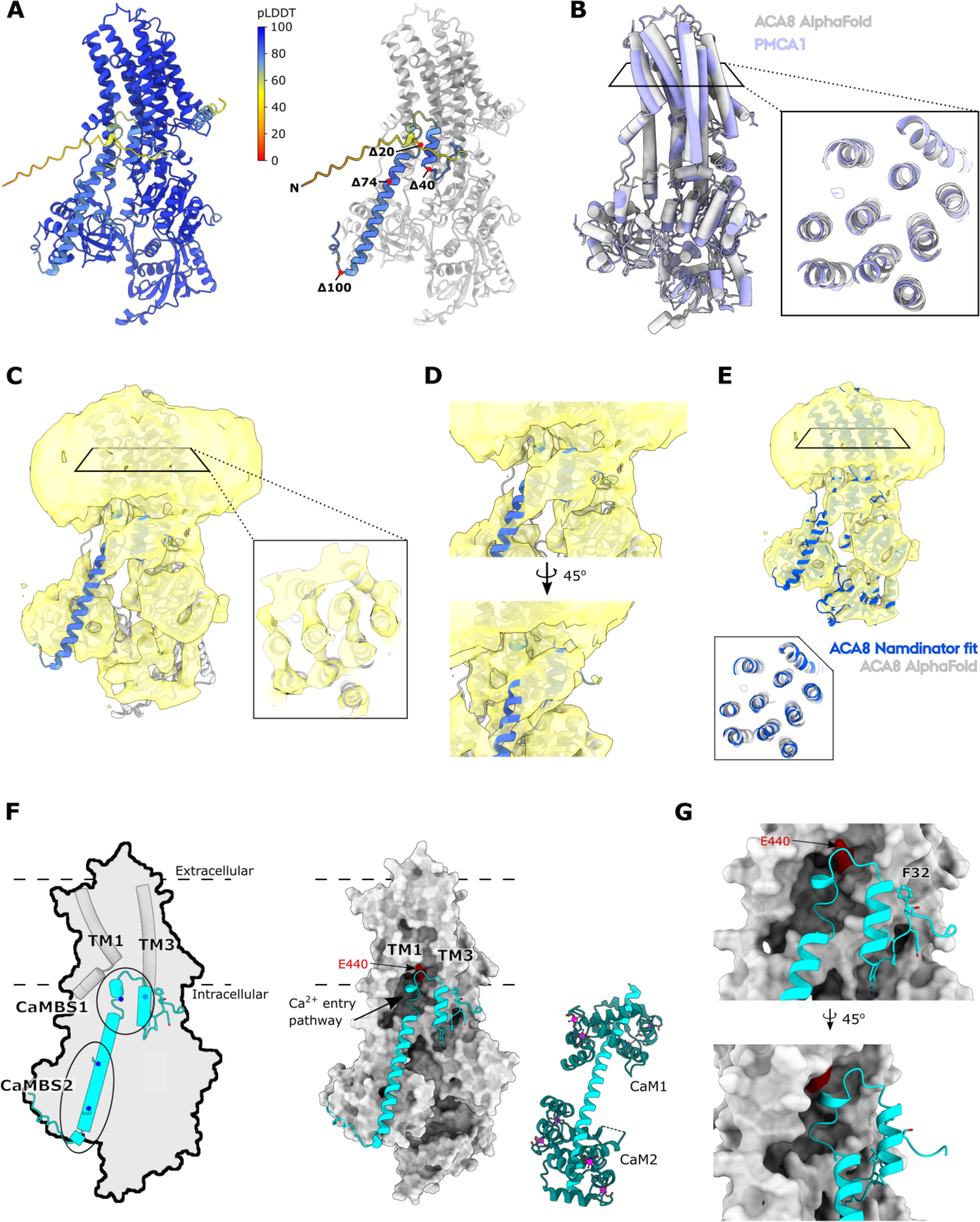
AlphaFold predicted structure of ACA8 and low resolution cryo-EM density. **A)** *Left:* The AlphaFold structure of ACA8 represented as cartoon and colored by confidence level, pLDDT. *Right:* Autoinhibitory domain binding. The core of the pump in grey with the autoinhibitory domain colored by confidence level with the N terminus and positions of the Δ20, Δ40, Δ74 and Δ100 truncations indicated. **B)** The AlphaFold predicted structure of ACA8 (core pump with residues 1-112 excluded) in gray aligned with the EM structure of PMCA1 in an E1 state (PDB: **6A69**) in lilac [44]. The cutout shows a cross section of the TM domain. **C)** Low resolution density of ACA8 in the E1 state (yellow) with the core of ACA8 AlphaFold predicted model in gray and the autoinhibitory domain colored by confidence level. Map contour level 1.2. The cutout shows a cross section of the TM domain with clear density for all 10 helices. **D)** Close up view of the area with excess density compared to the core structure. The density fits with the position of the autoinhibitory domain in the AlphaFold predicted structure. Residues 1-26 left out for simplicity. Map contour level 1.2. **E)** Low resolution density of ACA8 in the E1 state (yellow) with the Namdinator fitted ACA8 predicted structure (Blue). The cutout shows a cross section of the TM domain. Map contour level 1.2. **F)** Autoinhibitory domain binding as predicted by AlphaFold and the low-resolution map. The core of ACA8 is gray and the autoinhibitory domain is cyan. Membrane position is indicated by the broken line. Residues 1-26 and 1047-1074 are left out for simplicity. *Left:* A schematic of autoinhibitory domain binding. TM1 and TM3 represented as tubes and the position of CaMBS1 and CaMBS2 on the autoinhibitory domain are indicated with the anchoring residues denoted by blue spots, the faded blue spot indicates the residue is positioned on the backside of the helix. *Right:* The core of the ACA8 structure represented as a surface with the autoinhibitory domain represented as a cartoon (cyan) side-by-side with the crystal structure of the autoinhibitory domain (cyan) with CaM bound (teal) (PDB: **4AQR**) [36]. The Ca^2+^ entry pathway of ACA8 is denoted by the arrow, E440 of the calcium binding site is indicated in red and TM1 and TM3 are indicated. **G)** Close up of the autoinhibitory domain binding in the Ca^2+^ entry pocket. The residues 31-35 are represented as sticks and E440 of the calcium binding site is indicated in red.

Importantly, the predicted structure matches a 7 Å resolution cryo-EM map of Δ20-ACA8 reconstituted in salipro and stabilized in the calcium-free E1 state with AMPPCP plus EGTA present (C). Despite a low resolution of the map, the AlphaFold predicted ACA8 structure adopting the E1 state clearly fits it (C), and excess density compared to the core of the pump overlaps with the AlphaFold predicted structure for the autoinhibitory domain (Figure 3C and D). A small difference between the AlphaFold predicted ACA8 structure and the cryo-EM E1 structure of ACA8 is the position of the N domain, which is a bit closer to the A and P domains in the cryo-EM structure than for the AlphaFold predicted ACA8 structure. Using Namdinator [45], a program that in a flexible manner fits template models into density maps, the N-domain of the AlphaFold predicted ACA8 structure is repositioned in the density, while the TM helices maintain the E1 conformation (Figure 3E). The autoinhibited state described here therefore represents an E1 state preceding calcium binding and autophosphorylation.

The agreement between the AlphaFold prediction for the autoinhibitory domain binding and the unbiased, low resolution 3D map invites further analysis of autoinhibition.

The autoinhibitory domain is predicted to bind at the Ca^2+^ entry pathway, near transmembrane helix 1 (TM1) and TM3, and with an extended helix approaching the A domain (Figure 3F). The extended helical part resembles the crystal structure of the autoinhibitory domain in complex with CaM, where the domain with two CaM bound adopts a fully extended helix [36] (Figure 3F).

In the full-length AlphaFold predicted ACA8 structure, the residues from 30 to 65 form a helical structure near the membrane, with residues 52-65 of CaMBS1 obstructing entry to the Ca^2+^ binding site (Figure 3F). Consistently, residues 30-35 that from our mutational data were shown to be important for autoinhibition form interactions near TM3 and are positioned next to a part of CaMBS1, which is predicted to be helical (Figure 3G). F32 is a core element of this interaction at TM3. Interestingly, residues 66-100 encompassing the second CaM binding site (CaMBS2) are more exposed and predicted as an extended helix that spans from the Ca^2+^ entry pathway to the A domain (Figure 3F).

## Discussion

Activity studies of truncated versions of ACA8 implicate the region before the first CaM binding site in autoinhibition. Removing the first 40 residues from ACA8, partial loss of autoinhibition is observed, while a Δ20 construct exhibits maximal autoinhibition, even compared to WT. Possibly, the first 20 residues form a highly charged and disordered tail (Figure S1) that increases the “solubility” of the autoinhibitory domain and facilitates the dissociation of the autoinhibitory domain more than the Δ20 construct, which therefore shows stronger autoinhibition. The stretch from residues 20 to 40 includes conserved motifs and may impact stability of the autoinhibitory interaction before the first CaM binding site. This makes a Δ40 construct resemble the Δ74-ACA8 construct with a high basal activity and only partial stimulation by CaM [36]. Still, yeast complementation studies *in vivo* show a difference between Δ40 and Δ74, where Δ74 allows for better growth (less autoinhibition) than Δ40.

The *in vivo* experiments revealed a particular importance of residues 30-35 and pinpointed the highly conserved phenylalanine, F32, which is essential for maintaining tight regulation (Figure 1E). This segment and specifically F32 could be an important area of interaction for the autoinhibitory domain with the core of the pump. Indeed, the AlphaFold prediction of the autoinhibited state, which is consistent with a low-resolution map for the E1 state of ACA8, shows the 30-35 segment together with CaMBS1 forming an intimate interaction with F32 at the core that restricts movements of TM3 and blocks the Ca^2+^ entry pathway of ACA8 (Figure 3F and 3G).

Consistent with the role of the entire 20-35 segment, several serine residues in the 20-30 segment can be phosphorylated and regulate the activity and CaM stimulation of the pump [46]. While Δ20 shows WT behavior *in vitro* and *in vivo*, Δ30 is deregulated in membrane activity assays, although still sufficiently autoinhibited that it will not complement Ca^2+^ transport *in vivo*.

The processive deregulation observed for N-terminal truncation constructs merited further in-depth investigation. Purified ACA8 samples in WT, Δ20 and Δ100 forms could be solubilized and purified with activity maintained for all three forms, and with autoinhibition maintained for WT and Δ20 (Figure 2A), consistent with membrane-based assays. Purified protein allowed for direct studies of autoinhibition and CaM activation, and for studies of the impact of specific lipids. Indeed, acidic phospholipids were found to activate and overrule CaM sensitivity of purified ACA8 (Figure 2A), in agreement with previous studies [39].

Purified ACA8 reconstituted in salipro lipid nanodiscs exhibited autoinhibition and allows for structural studies of the autoinhibited state. The cryo-EM sample was favorable for the Ca^2+^-free E2-BeFx complex mimicking the outward-open E2P state, and the structure was determined at 3.3 Å resolution overall. ACA8 shows the expected P-type ATPase features and is similar to the E2-BeFx complex of SERCA (Figure 2E). Unlike for P4-ATPases [43], however, the E2P state of ACA8 did not reveal any density for the autoinhibitory domain, and consistently, Saffioti et al. also showed that the autoinhibitory interaction is disrupted in the E2P state [47].

However, the AlphaFold predicted structure with the autoinhibitory domain bound indicates an E1 state, and a low resolution cryo-EM structure obtained for ACA8 in the E1 state was consistent and showed overlapping density near the membrane by TM3 (Figure 3C and D). The predicted mode of binding for the autoinhibitory domain in the Ca^2+^ entry pathway would not be possible in the E2P state of ACA8, where TM1 is closed in on TM3 effectively obstructing Ca^2+^ release/entry in that direction and also autoinhibitory domain binding (Figure 4A and B). Oppositely, binding of the autoinhibitory domain mimics the cytoplasmic occlusion of the Ca^2+^ entry pathway that occurs in the E2P state, so autoinhibition and the E2P state are mutually exclusive. Autoinhibition depends on a flexible binding of the autoinhibitory domain, and a low basal activity is observed, in particular for the detergent solubilized form.

**Figure 4:**
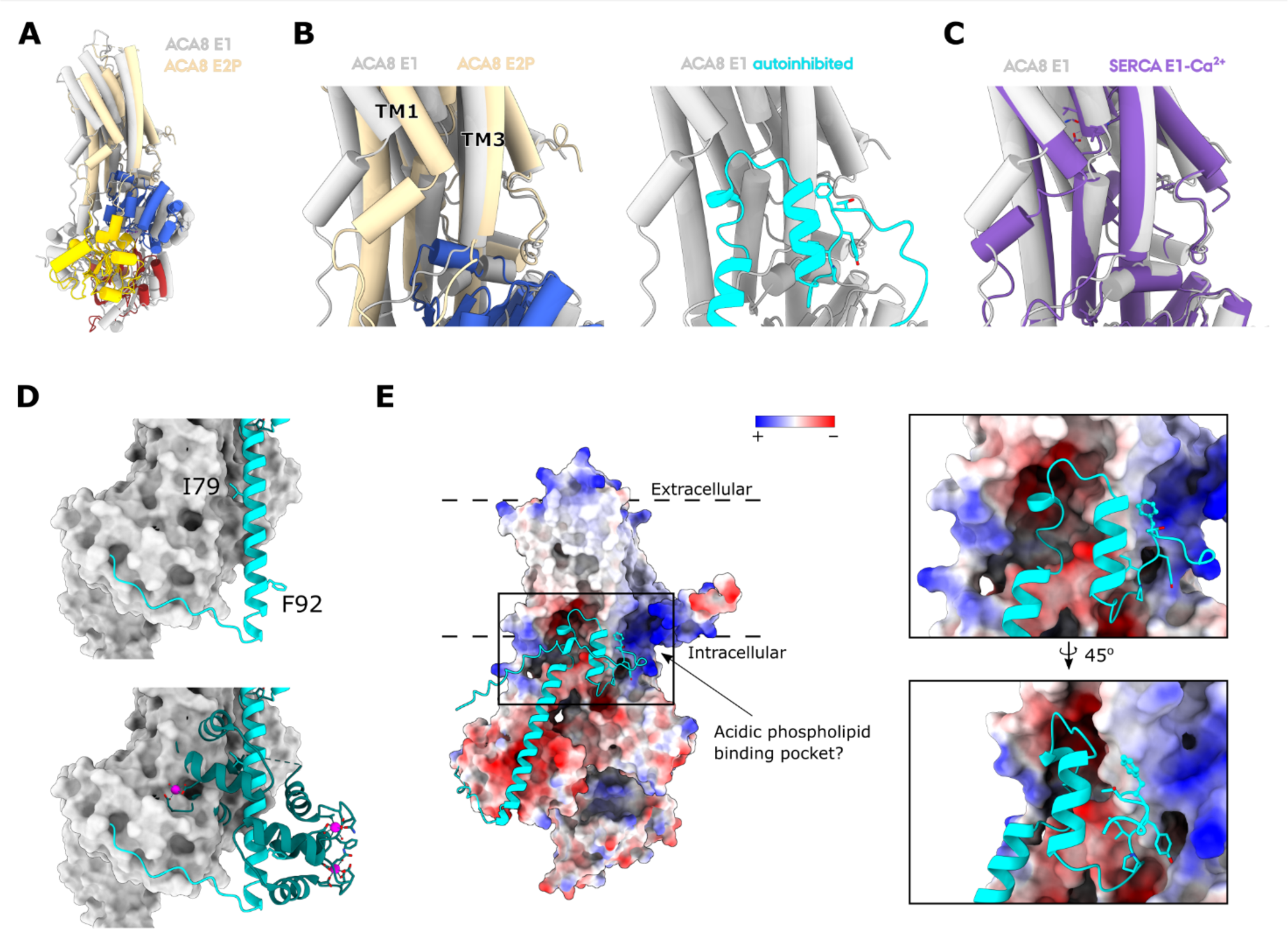
ACA8 structure comparison and activation model. The core of the ACA8 E1 state predicted by AlphaFold and the low-resolution density in gray aligned with ACA8 E2P cryo-EM structure colored by domain (TM domain in wheat, A domain in yellow, P domain in blue and N domain in red), represented as cartoon with tubular helices. The autoinhibitory domain, residues 1-112, of the E1 state is excluded for clarity. Alignment on TM7-10 segment. **A**) Zoom in on the Ca^2+^ entry pathway of ACA8 with a side-by-side view of E1 and E2P alignment and autoinhibitory domain binding. *Left:* Comparison of the Ca^2+^ entry pathway for the E1 and E2P state of ACA8. TM1 is shifted inward towards TM3 in the E2P structure compared to E1 thereby blocking Ca^2+^ entry. *Right:* AlphaFold predicted E1 structure with the autoinhibitory domain (cyan) blocking Ca^2+^ entry. **C)** Zoom in on the Ca^2+^ entry pathway of ACA8 aligned with the E1 Ca^2+^ AMPPCP SERCA structure (PDB: **1T5S**) **D)** The core of the AlphaFold predicted structure of ACA8 as a grey surface and the autoinhibitory domain in cyan. *Top*: Close up side view of the elongated helix with CaMBS2. The anchoring residues exposed and available for CaM interaction are shown as sticks. *Bottom*: CaM binding to the exposed CaMBS2. CaM in teal with Ca^2+^ in magenta. CaMBS2 of the AlphaFold structure was aligned with CaMBS2 of the crystal structure of the autoinhibitory domain of ACA8 with two CaM bound (PDB: **4AQR**) (Tidow et al., 2012). **E)** The core of the Alphafold predicted structure represented as an electrostatic surface, red negatively charged, white neutral and blue positively charged, with the autoinhibitory domain is in cyan and the potential acidic phospholipid site is indicated by an arrow. The cutouts show a close up view of the potential acidic phospholipid binding site near F32.

The AlphaFold predicted structure furthermore showed CaMBS2 being exposed as an extended helix, which was partially corroborated by density for the low resolution E1 structure (Figure 3C and D). Likely, it forms a very flexible interaction, and it might be difficult to obtain a high-resolution structure for the autoinhibited state. The autoinhibitory domain was also lacking in a 4.1Å resolution cryo-EM structure of mammalian PMCA1 in the E1 state [44].

The predicted mode of autoinhibition locks the protein in a pre-activated state, where flexibility of the autoinhibitory domain allows rare incidents of Ca^2+^ entry and basal activity, or full activation by CaM binding distracting CaM binding sites from autoinhibitory interactions. Locking the protein in the E1 state makes functional sense, as the pump is then ready to start transport as soon as autoinhibition is relieved or Ca^2+^ enters the binding site, and the entry pathway closes upon Ca^2+^ binding/occlusion (Figure 4C).

Similary, autoinhibition of the P4-ATPase lipid flippases also traps the state immediately preceding transport, in this case an autoinhibited E2P. In the free E2P state, P4-ATPases are open towards the extracellular side of the membrane ready to bind and flip a lipid molecule as soon as autoinhibition is lifted [43].

The autoinhibited state as predicted by AlphaFold (that we consider validated by the clear match to the low resolution cryo-EM structure for the E1 state of ACA8) suggests that futile ATPase cycles are prevented by separation of the ATP binding site and the reactive aspartate side chain (Figure S12). However, modulatory ATP binding [48] would be allowed and could contribute also to the balance between autoinhibition and activity. However, the resolution of the map for the E1 state we present here is too low to observe nucleotide binding.

The mapping of the autoinhibitory domain also explains what we observe here and that has been observed in previous biochemical data, namely that there is a difference in activation by CaM for Δ74 and Δ100, with Δ74 losing autoinhibition but still showing some activation by CaM [36]. The Δ74 construct would retain a helical segment associating with the A domain that potentially impairs the approach of the A domain towards the P domain in the E1 to E1P transition (Figure S13). Binding of CaM to CaMBS2 would dislocate the helix and leave a fully active pump with transition between E1 and E2 no longer being hampered.

Previously Tidow et al. proposed a two-step model for activation by CaM where CaM binding first occur at CaMBS1 followed by binding to CaMBS2 [36]. Based on isolated peptides, biophysical data showed CaMBS1 as a high-affinity binding site and CaMBS2 as a lower affinity site [36]. However, from the E1 autoinhibited model presented here, we observe that CaMBS1 is closely bound near the Ca^2+^ entry site and near the membrane (Figure 3F and G), whereas CaMBS2 is far more exposed. The two anchoring residues of CaMBS2, I79 and F92 [36] are exposed, in particular F92, and binding of CaM to CaMBS2 would effectively result in a sterical clash between CaM and the A domain, destabilizing the autoinhibited state (Figure 4D). This suggests that CaMBS2 is the first site to bind CaM, followed by CaMBS1. Binding of CaM to the exposed CaMBS2 would force at least a partial dislocation of the autoinhibitory domain that could translate to CaMBS1 becoming then more accessible to CaM binding.

However, ACA8 is also highly activated by acidic phospholipids and looking at an electrostatic surface representation for the AlphaFold predicted structure of ACA8 there is a clear positively charged pocket at the membrane interface (Figure 4E). This potential lipid binding pocket is positioned near the area where F32 interacts with the core of the pump, and binding of an acidic phospholipid could potentially directly impact autoinhibitory domain anchoring resulting in CaMBS1 being dislodged first.

Our observation on ACA8 activity and the model for the autoinhibited E1 state propose a model for ACA8 regulation, where the 20-40 segment preceding CaMBS1 is important for anchoring of the autoinhibitory domain. Destabilizing or truncating the anchoring segment favors dislocation of the autoinhibitory domain from the Ca^2+^ entry pathway. Upon CaM activation, CaM might first bind to CaMBS2 of ACA8 thereby promoting further dislocation of the autoinhibitory domain and binding of the second CaM at CaMBS1 to relieve blockade of the Ca^2+^ entry pathway and provide full activation (Figure 5). The binding of the autoinhibitory domain can also be directly disrupted by the presence of acidic phospholipids and potentially by phosphorylations at the 20-30 segment – an effect we also see by mutation of conserved F32 to an alanine (Figure 5). Potentially, two routes of ACA8 activation therefore exist, and this is reminiscent of the two routes observed also for mammalian PMCA, namely through CaM with neutral lipids and through aSN with negatively charged lipids, respectively.

**Figure 5:**
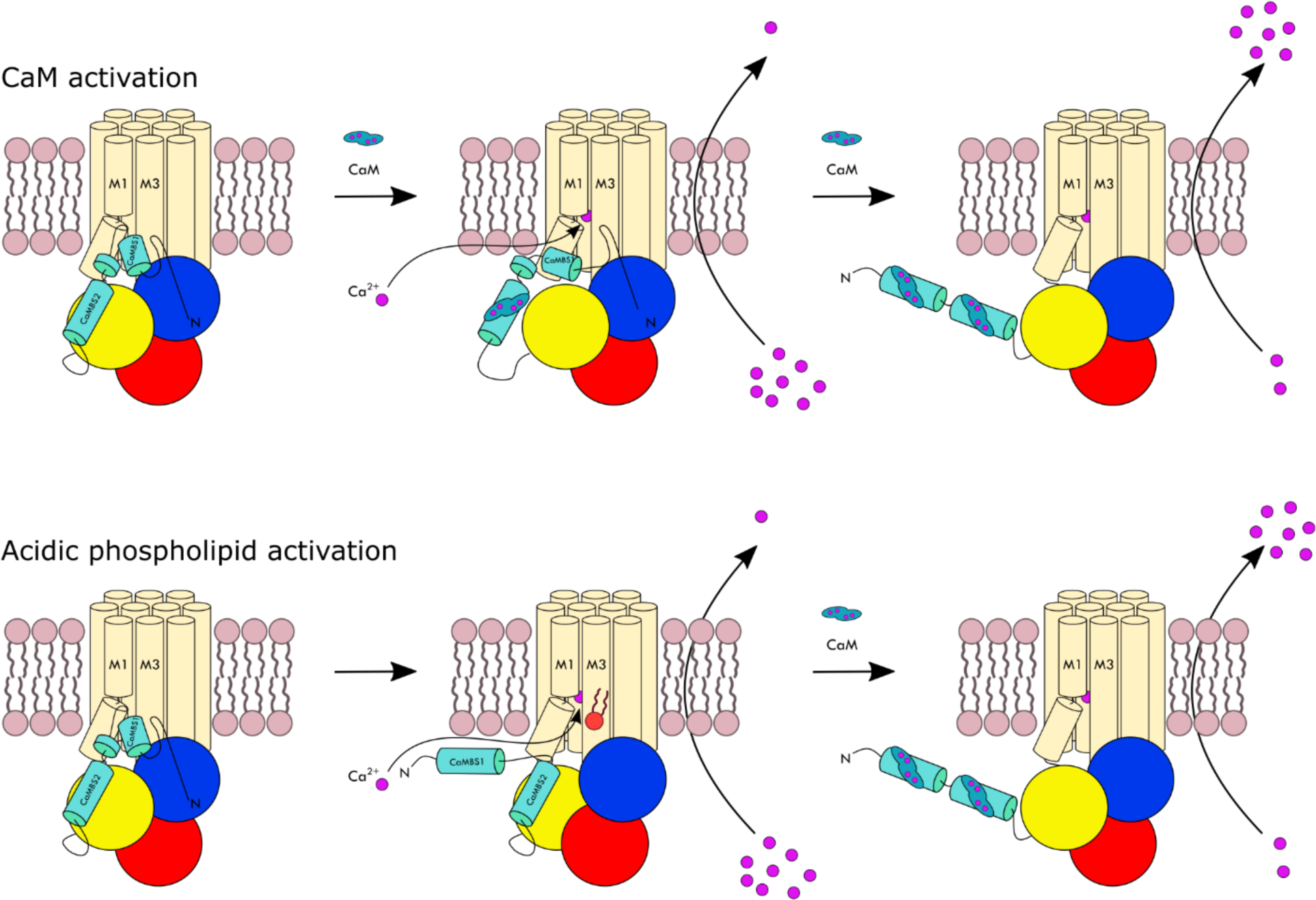
Model for displacement of the autoinhibitory domain of ACA8. An illustration of the proposed model for activation of ACA8. The TM domain in wheat, the A domain in yellow, the P domain in blue and the N domain in red. The autoinhibitory domain in cyan, with CaM in teal and Ca2+ in magenta Top: Activation by CaM. The autoinhibitory domain is bound near TM3 with the CaMBS1 blocking Ca^2+^ entry. Binding of CaM to CaMBS2 destabilizes the binding of CaMBS1 at the Ca^2+^ entry pathway exposing it further for CaM binding. CaM binding at CaMBS1 fully relieves autoinhibition. Bottom: Activation by acidic phospholipids. The presence of acidic phospholipids disrupts the interaction between CaMBS1 and the Ca^2+^ entry pathway and TM3 displacing CaMBS1 resulting in an activated pump that can transport Ca2+ across the membrane. Binding of CaM fully relieves autoinhibition and ACA8 is fully active and can rapidly transport Ca^2+^ across the membrane.

## Materials and Methods

### Construct formation

The Δ74- and Δ100-ACA8 constructs were previously described [36] and expression plasmids were supplied by Michael Palmgren. The other N-terminally truncated ACA8 constructs (Δ20-, Δ30-, Δ35-, Δ37-, Δ40-ACA8) were cloned into a pYES2 vector with a GAL1 promoter and ura3 gene for selection. The Δ20- and Δ30-ACA8 constructs were cloned using ligation dependent cloning whereas Δ35-, and Δ37-ACA8 were cloned using homologous recombination (pYES2 vector linearized by BamHI and XhoI treatment was mixed with PCR product of the construct containing the right overlap sequences, ratio 1:15) followed by yeast miniprep and up-formation in *E. coli*. The Δ40-ACA8 construct was created by amplification of the entire vector except the sequence encoding for the first 40 residues using primers with complementary overhangs. The Δ20-, Δ30-, Δ35-, Δ37-, Δ40-ACA8 constructs all have an N-terminal (His)10-tag.

The F32A mutants of WT and Δ20-ACA8 were created using a QuikChange site directed mutagenesis kit (Agilent technologies) and the following primers:

F32A forward primer: GACACT**GCC**TACATCCCATCGAAGAATGCTTC

F32A revers primer: GATGTA**GGC**AGTGTCGCTATCAGAGTCTGCGT

### Yeast transformation and cell growth

The ACA8 constructs were expressed in the calcium pump deficient *Saccharomyces cerevisiae* strain K616 (MATa pmr1::HIS3 pmc1::TRP1 cnb1::LEU2, ade2, ura3). Transformations were performed according to the lithium acetate, single-stranded DNA, PEG method [49]. Transformed cells were selected on uracil deficient plates, picked and grown in minimal media (2 % glucose, 0.7 % yeast nitrogen base, 0.8% Complete Supplement Mixture -URA, 40 mg/L adenine, 10 mM CaCl2) at 30 °C. Cells were induced in rich galactose media (2 % galactose, 2 % Bactopeptone, 1 % yeast extract, 40 mg/L adenine, 10 mM CaCl2) and incubated for 16 hours at 30 °C. The cells were harvested by centrifugation and resuspended in GTEB20 buffer (20 % glycerol, 50 mM Tris-HCl pH 7.6, 1 mM EDTA, 5 mM βME, Complete^TM^ protease inhibitor).

### Yeast complementation assay

The yeast complementation assay was performed as earlier described [20]. Briefly, transformed K616 cells were taken straight from the Uracil selection plate, suspended in water and dropped out on plates containing either 2% glucose and 10 mM EGTA, 2% glucose and 10mM CaCl2 (SDA plates), 2% galactose and 10mM EGTA, or 2% galactose and 10 mM CaCl2 (SGA plates). Two independent experiments were performed.

It has previously been shown that only K616 cells expressing an active Ca^2+^-ATPase can complement the K616 yeast strain, which is a strain with knock-out of the two endogenous Ca^2+^-ATPases, PMR1 and PMC1 [20, 37]. K616 cells need an active Ca^2+^ pump to grow on CaCl2 depleted media, and it is believed that the activity maintains vital, internal Ca^2+^ stores [37]. Cells expressing an autoinhibited ACA8 therefore will not be able to grow on CaCl2 depleted media, while cells expressing a constitutively active ACA8 do. The constitutively active Δ100-ACA8 construct has previously been shown to complement the K616 strain [36] and was used as a positive control for growth on CaCl2 depleted media.

### Yeast membrane isolation

Cells were cracked with glass beads in a planetary mono mill, “pulverisette 6” (Fritsch). Cell debris was pelleted at 1000 x g for 20 min. The supernatant was centrifuged 20 min at 20,000 x g and the subsequent supernatant was ultracentrifuged for 75 min at 220,000 x g, pelleting the ACA8-containing membranes. The pellet was resuspended in a high salt GTEB20 (1M KCl) buffer to remove peripheral membrane proteins and subsequently repelleted by ultracentrifugation. The washed membranes were resuspended in GTEB20, flash frozen in liquid nitrogen and stored at -80 °C until activity testing or purification. The membrane protein concentration was determined using a Bradford assay.

### Ca^2+^ ATPase activity assays on membrane bound ACA8

400µL reactions with 4 *μ*g of membrane protein samples containing the different constructs were assayed in four different buffers: 50 mM KNO3, 5 mM NaN3, 0.25 mM Na Molybdate, 5 mM (NH4)2SO4, 40 mM Bis Tris Hepes pH 7.5, 0.1 mg/mL Brij58, 1 *μ*M A23187, 3 mM MgSO4, 1 mM EGTA with either i) 1.125 mM CaCl2 (corresponding to 35 *μ*M free Ca^2+^, estimated using the program WEBMAXCLITE v1.15), ii) 500 nM CaM, iii) 500 nM CaM and 1.125 mM CaCl2, or iv) completely without CaM and CaCl2. Full activation of membrane bound WT ACA8 is observed with addition of 0.5 µM CaM (Figure S14).

The ATPase activity was tested by a time course assay at 30 °C monitoring the generation of free inorganic phosphate in a modified calorimetric assay adapted from Baginski et al. [50]. At specific time points over the course of an hour the reactions were stopped and developed for absorption spectroscopy at 860 nm. The Ca^2+^ dependent activity was calculated by subtracting activities without Ca^2+^ from activities with Ca^2+^. The activity of ACA8 with only CaM and no Ca^2+^, did not show any additional activity compared to ACA8 without both CaM and Ca^2+^. The fold activation by CaM, found by dividing the activity with CaM present with the basal activity, was determined from biological duplicates all tested in technical triplicates or more. The CaM dependent activation was determined from 6 (Δ20-, Δ40-, Δ74-, Δ100-ACA8, WT F32A ACA8, Δ20 F32A ACA8) or 12 (Δ30-, Δ35-, Δ37-ACA8) different experiments.

### CaM expression and purification

Calmodulin 7 (CaM7) from *A. thaliana* was used for all experiments and was recombinantly expressed in *E. coli* and purified with an N-terminal (His)6-tag. Chemically competent C41 *E. coli* cells were transformed with a pET15b CaM7 plasmid. Cells were grown at 37 °C, induced by 1 mM IPTG at an OD600 of 0.6, and growth continued for 16 hours at 21 °C. Cells were harvested by centrifugation and lysed using sonication in buffer (40 mM Tris, 300 mM NaCl, 5 mM CaCl2, 15 mM imidazole, Complete^TM^ protease inhibitor). The lysate was centrifuged at 20,000g for 1 hour. CaM7 was purified using a standard procedure using an immobilized Ni^2+^-column (HisTrap, GE Healthcare). Load buffer (40 mM Tris, 300 mM NaCl, 5 mM CaCl2) was used for loading and washing of the column, and the pure protein eluted in an elution buffer (40 mM Tris, 300 mM, NaCl, 5 mM CaCl2 and 500 mM imidazole).

### Sequence alignment

The sequence alignment was produced using CLUSTAL omega [51]. The sequence alignment was performed on 8 different ACAs from *A. thaliana*, all found as annotated entries in Uniprot. The alignment was analysed using Jalview [38]. Uniprot accession codes: Q9LF79 (ACA8), O81108 (ACA2), O22218 (ACA4), Q37145 (ACA1), Q9SZR1 (ACA10), Q9LU41 (ACA9), Q9M2L4 (ACA11), O64806 (ACA7).

### ACA8 purification

The WT, Δ20- and Δ100-ACA8 were purified based on modified protocols from previously published procedures for Ca^2+^-ATPases [52]. ACA8-containing yeast membranes with a total membrane protein concentration of 5 mg/mL were solubilised in solubilisation buffer (50 mM Tris-HCl pH 7.6, 100 mM KCl, 20% glycerol, 3 mM MgCl2, 5 mM β-mercaptoethanol (βME), 1 mM PMSF, 1 µg/mL chymostatin, 1 µg/mL pepstatin A, 1 µg/mL leupeptin) with 7.5 mg/mL DDM (dodecyl maltoside) for 90 min at 4 °C.

Solubilised membranes were ultracentrifuged at 200,000 x g to pellet unsolubilised membranes. The supernatant was filtered (0.45µm) and loaded onto a Ni^2+^-affinity column (HisTrap FF, GE Healthcare 5 mL column) equilibrated in low imidazole buffer (50 mM Tris-HCl pH 7.5, 10 mM imidazole, 200 mM KCl, 10 % glycerol, 3 mM MgCl2, 5 mM βME, 0.2 mM LMNG (Lauryl Maltose Neopentyl Glycol)). The column was washed in the low imidazole buffer.

For WT- and Δ20-ACA8 1.5 column volume (CV) of buffer with 200 units of thrombin was loaded onto the column and left overnight for cleavage at 4 °C. Cleaved WT- and Δ20-ACA8 was eluted from the column in low imidazole buffer.

For Δ100-ACA8 the protein was eluted off the Ni^2+^ column in high imidazole buffer (50 mM Tris-HCl pH 7.5, 500 mM imidazole, 200 mM KCl, 10 % glycerol, 3 mM MgCl2, 5 mM βME, 0.2 mM) and dialysed in buffer without imidazole and cleaved with TEV overnight. The cleaved Δ100 ACA8 was then reapplied to the Ni^2+^ column and collected as flow-through from the Ni^2+^-affinity column equilibrated in the low imidazole buffer. The ACA8 was relipidated with 5 µg DOPC (dioleoyl phosphatidylcholine) per mL of eluted ACA8 (DOPC stock: 5 mg/mL DOPC in 10 mg/mL LMNG) and subjected to size exclusion chromatography (SEC) on a superose 6 column in a SEC buffer (50 mM Tris-HCl pH 7.5, 200 mM KCl, 3 mM MgCl2, 10% glycerol, 0.02 mM LMNG, 5 mM βME).

### Salipro reconstitution of Δ20-ACA8

Nickel affinity purification was performed for Δ20-ACA8 as described in the “ACA8 purification”-section above, and the protein was mixed with detergent solubilized SBPC (soybean phosphatidylcholine from Avanti polar lipids) using a molar ratio of 1:80 and incubated for 30 minutes at room temperature (RT). Saposin A was added to a molar ratio of 1:25 and the sample was incubated for an additional 10 min at RT. Lastly, the sample was diluted below the critical micelle concentration (CMC) of DDM to 0.0055% in dilution buffer (200 mM KCl, 50 mM Tris-HCl pH 7.5). To optimize the amount of reconstituted protein, multiple small-scale reconstitution reactions (100 µL prior to dilution) were performed. The small-scale reconstitution reactions were pooled after dilution and concentrated using a 100kDa MWCO Vivaspin protein concentrator (GE Healthcare). The sample was subjected to size exclusion chromatography (SEC) on a Superdex 200 increase 10/300 GL column (GE Healthcare) in SAP-buffer (200 mM KCl, 50 mM Tris-HCl pH 7.5).

### Ca^2+^ ATPase activity assays on purified ACA8

Activity assays on purified protein were performed using 1 or 2 µg protein in 400 µL reaction buffer (0.125 µg and 0.25 µg per 50 µL reaction) (approx. 0.04 µM and 0.08 µM) with 50 mM KNO3, 5 mM NaN3, 0.25 mM NaMob, 40 mM Bis Tris Hepes pH 7.2, 3 mM MgSO4, 1 mM EGTA. Activity was tested i) without CaCl2, ii) with 1.125 mM CaCl2 (corresponding to 35 *μ*M free Ca^2+^), or iii) with 830 nM CaM and 1.125 mM CaCl2.

Prior to the reaction the purified protein was incubated with 5 µg or 10 µg phosphatidyl choline lipids from soy bean (1:5 w/w protein to lipid ratio). The ATPase activity was tested by a time course assay at 30°C as described for the membrane bound protein over 30 minutes. For the salipro reconstituted Δ20-ACA8 the protein was not incubated with lipid prior to the reaction since lipids were present in the salipro disc.

### Cryo-EM grid preparation

C-flat holey carbon, CF-1.2/1.3 300-mesh Cu (Protochips) (data for E2P state) or CF-2/2 300-mesh Cu (data for E1 state) grids were glow discharged for 45 s at 15 mA using the PELCO easiGlow. Salipro-reconstituted Δ20-ACA8 was applied at a final concentration of 0.67 mg/mL with an addition of 1 mM EGTA, 1 mM BeFx and 3 mM MgCl2 to stabilize an E2-BeFx form mimicking the E2P conformation or at a final concentration of 0.6 mg/mL with 1 mM AMPPCP, 1 mM EGTA, 3 mM MgCl2 to mimic the E1 conformation. The grids were plunge-frozen in liquid ethane after blotting for 4 s with the sensor blotting setting using the Leica GP2 plunge freezer operating at 10 °C and 90% humidity. Grids were stored in liquid nitrogen.

### Cryo-EM data acquisition

Electron micrographs were collected at 300 kV with a Titan Krios G3i (Thermo Scientific) equipped with a Gatan K3 camera operating in super resolution mode. For the E2P data the nominal magnification was 130,000x corresponding to a pixel size of 0.647 Å. A total of 5297 movies were recorded automatically with EPU software (Thermo Scientific) at a total electron dose of 60.1 e/Å^2^. For the E1 data the nominal magnification was 130,000x corresponding to a pixel size of 0.66 Å. Two datasets were collected with a total of 5988 movies (dataset 1) or 5050 movies (dataset 2) and were recorded automatically with EPU software (Thermo Scientific) at a total electron dose of 62.2 e/Å^2^ (dataset 1) or 61.8 e/Å^2^ (dataset 2). Cryo-EM data acquisition statistics can be found in Table S2.

### Cryo-EM data processing

#### Higher resolution E2P data

Movies were motion-corrected, and contrast transfer function (CTF) was estimated using cryoSPARC2 [53], and 941,952 particles were automatically picked using a circular blob. Particles were initially extracted in a 96-pixel box size (2.59 Å/pixel). 2D classification was run on the particles, and *ab initio* references were created from a subset of 100,000 particles. A representative micrograph (Figure S15A) and 2D classes (Figure S15B) are shown in supplemental information.

One protein-like reference and multiple junk references from *ab initio* classification were used in two rounds of heterogeneous refinement of the 941,952 particles to sort out junk particles in 3D, resulting in 346,176 particles. Selected particles were re-extracted in a 416-pixel box Fourier-cropped to 256-pixel box (1.08 Å/pixel) before 2 more rounds of heterogeneous refinement followed by non-uniform refinement resulting in 193,419 particles and a resolution of 3.33 Å of the resulting map. Finally, the particle stack was CTF-refined using Local CTF refinement and locally refined using a mask masking out the Salipro disc of the volume resulting in a map of the protein with a resolution of 3.29 Å. FSC curve (Figure S15C) and distribution of particle orientations (Figure S15D) are shown in supplemental information. Cryo-EM data processing statistics can be found in Table S2.

#### Low-resolution E1 data

Movies were motion corrected as for the E2P dataset and 996,849 particles from dataset 1 and 752,205 particles from dataset 2 were automatically picked.

First, a protein class was obtained from *ab initio* modelling of the best 2D classes followed by homogeneous refinement. 3 rounds of heterogeneous refinements were performed with 2 junk classes and the protein class. The heterogeneous refinements were followed by *ab initio* modelling performed with 7 Å as maximum resolution and 3 classes. The particles from the best *ab initio* model was followed by homogeneous refinement. A final *ab initio* run with a maximum resolution of 5 Å and 3 classes were followed by homogeneous refinement on the 40,686 particles from the best *ab initio* model. Lastly a non-uniform refinement was performed yielding a final model with overall resolution of about 7 Å. Representative 2D classes (Figure S16A), FSC curve (Figure S16B), particle orientation distribution (Figure S16C) and Cryo-EM data processing statistics (Table S2) can be found in supplemental information.

### Model building and refinement

An initial model of ACA8 was created using Modeller [54] and the structure of SERCA in an E2P state. The initial model was fit in the density using Namdinator [45] then manually adjusted in *Coot* [55] using ModelAngelo [56] to aid model building.

Several rounds of real space refinement in Phenix [57] followed by manual adjustments in *Coot* were performed. Validation statistics can be found in Table S3. Cross validation was performed determining FSCwork and FSCfree (Figure S17) [58]. Structural figures were prepared using ChimeraX [59].

### Structure prediction using AlphaFold

The AlphaFold structure predictions of the ACA8 was performed with a local installation of AlphaFold [60] using the AlphaFold multimer setting [61]. All output models were manually inspected and evaluated. To probe the validity of the predicted binding mode for ACA8 the structures of other ACA and close ACA8 Blast matches were also predicted. All models showed a similar predicted fold and mode of autoinhibitory domain binding as for the ACA8 model. Structures were represented using ChimeraX [59].

## Data Availability

The Cryo EM density map of ACA8 E2P structure is deposited in the Electron Microscopy Data Bank under accession number **EMD-18506** with the E1 map as an additional map. Coordinates for the ACA8 E2P model have been deposited in the protein Data Bank under accession number 8QMP.

## Acknowledgements

This study was supported by Lundbeck Foundation grant no. DANDRITE-R248-2016-2518 and grant no. R310-2018-3713 as well as The Danish National Research Foundation, Center for Membrane Pumps in Cells and Disease – PUMPkin and The Independent Research Fund Denmark – Natural Sciences, grant no. 7014-00328B.

We would like to acknowledge the EMBION Danish National Cryo-EM Facility for providing access to instruments and aid in data acquisition. We would like to thank Jesper L. Karlsen, Rasmus Kock Flygaard and Milena Timcenko for important inputs and guidance regarding data processing. We are grateful to Dr. Michael Habeck, Magnus Kjærgaard, and Dr. Florian Hilbers for valuable discussions and help.

## Author Contributions

STL, CFN and JDK performed the experiments. STL analysed the data. JDK and STL performed Cryo-EM data collection and processing. PN supervised the experiments and data analysis. LRP and MP aided in the design and initial set up of the complementation assay. STL wrote original draft preparation. STL, JDK and PN writing review and editing with input from all authors.

