## Supplementary material for "Conserved N-terminal Regulation of the ACA8 Calcium Pump with Two Calmodulin Binding Sites": Supplemenatry material

### Supplementary information

Includes figures S1 to S17 and tables S1 to S3

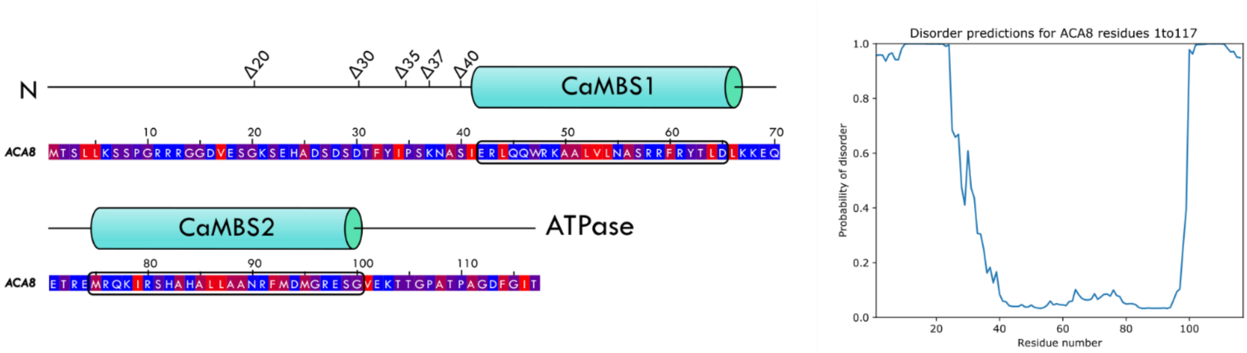

**Figure S1: Polarity and disorder of N terminus.**

*Left:* Polarity of the residues in the autoinhibitory domain of ACA8. Ranging from red (non-polar) to blue (polar). *Right:* Disorder prediction for the N-terminus and autoinhibitory domain of ACA8. Residues 1-25 is predicted to be highly disordered. Disorder prediction was performed using ODiNPred (Dass *et al*, 2020).

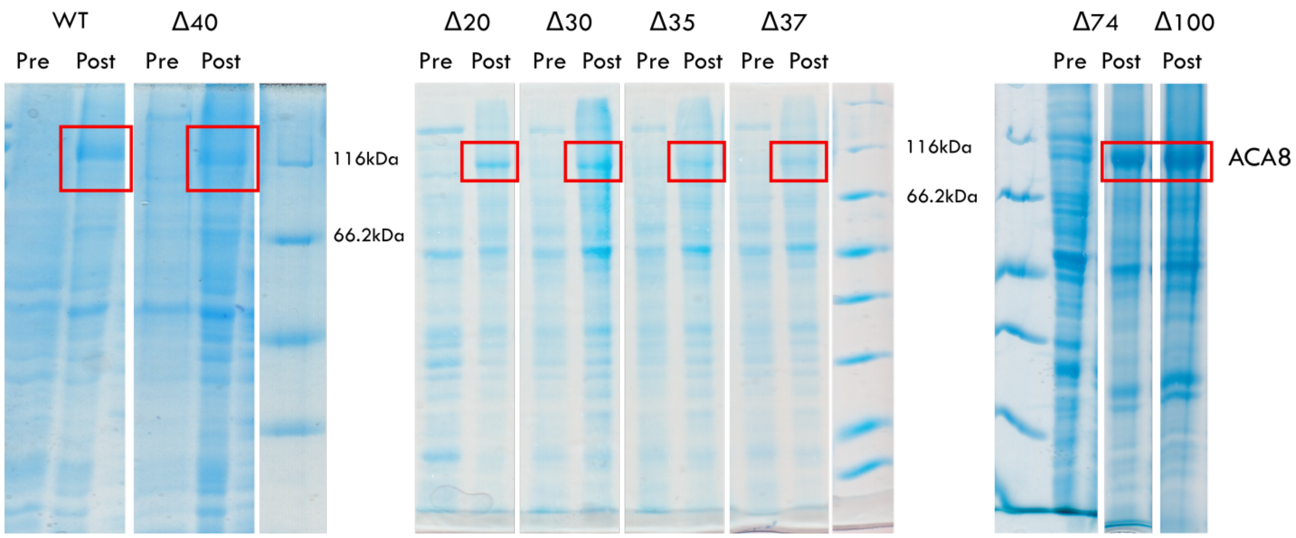

**Figure S2: Expression levels of truncation constructs.**

Coomassie stained SDS-PAGE gel of the isolated cell membranes, showing the expression of the ACA8 constructs. Pre and post induction samples for the constructs (only the Δ74 pre induction sample for Δ74 and Δ100 is part of the figure). The ACA8 band is marked by the red rectangles.

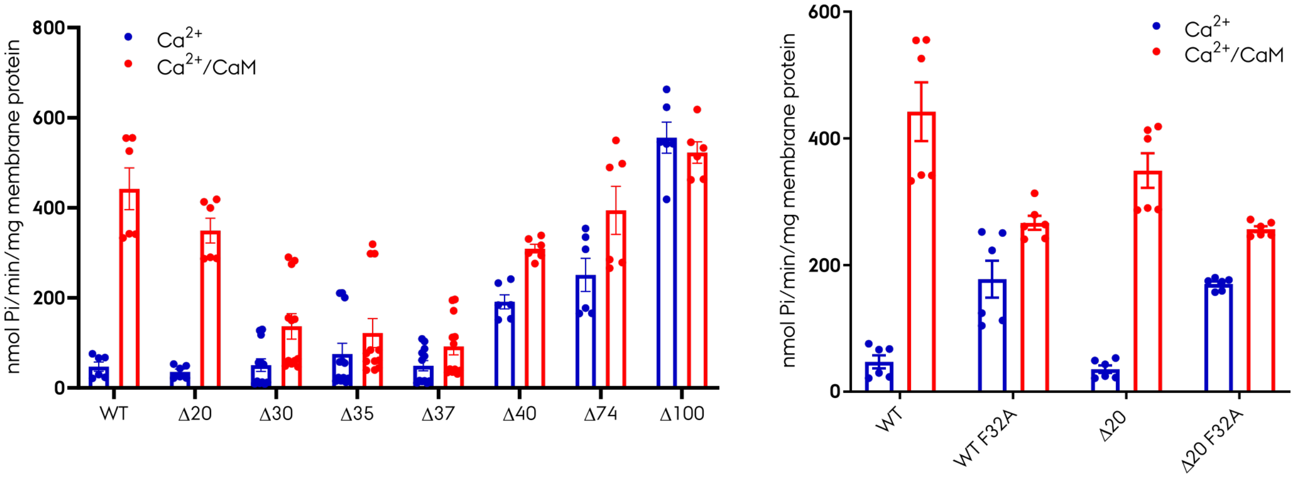

**Figure S3: Specific activities for ACA8 constructs.**

*Left:* The specific activity of the truncation constructs in the presence of $Ca^{2+}$ and $Ca^{2+}$/CaM. *Right:* The specific activity of the F32A mutants in the presence of $Ca^{2+}$ and $Ca^{2+}$/CaM.

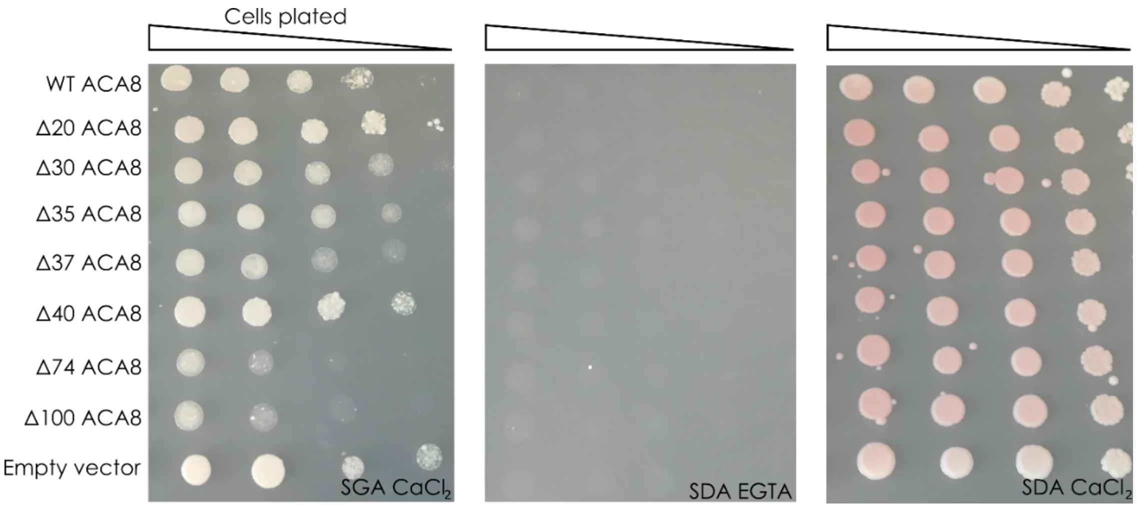

**Figure S4: Complementation assay – Control plates for truncation constructs.**

Galactose with $CaCl_2$ (SGA $CaCl_2$), Glucose without $CaCl_2$ (SDA EGTA) and Glucose with $CaCl_2$ (SDA $CaCl_2$).

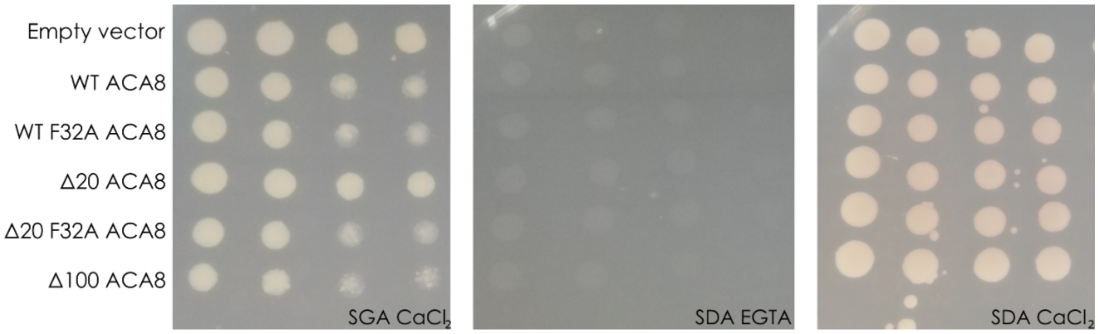

**Figure S5: Complementation assay – control plates for F32A mutants**.

Galactose with $CaCl_2$ (SGA $CaCl_2$), Glucose without $CaCl_2$ (SDA EGTA) and Glucose with $CaCl_2$ (SDA $CaCl_2$).

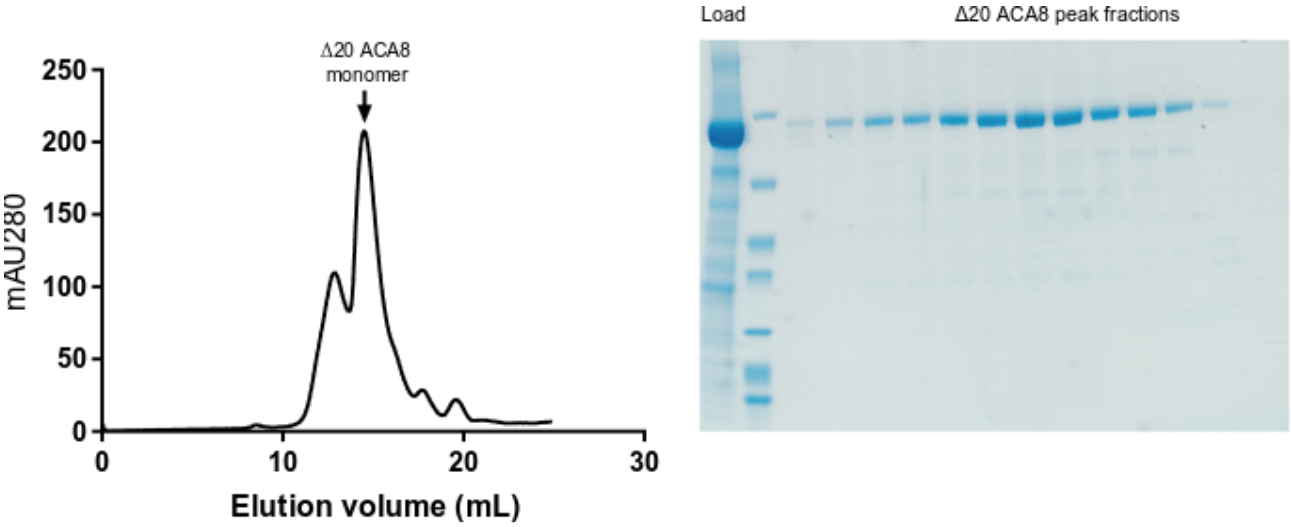

**Figure S6**: **SEC profile and SDS-PAGE from Δ20-ACA8 purification.**

*Left:* The elution profile for Δ20 with the monomer peak denoted by an arrow. *Right:* The SDS PAGE of the peak fraction.

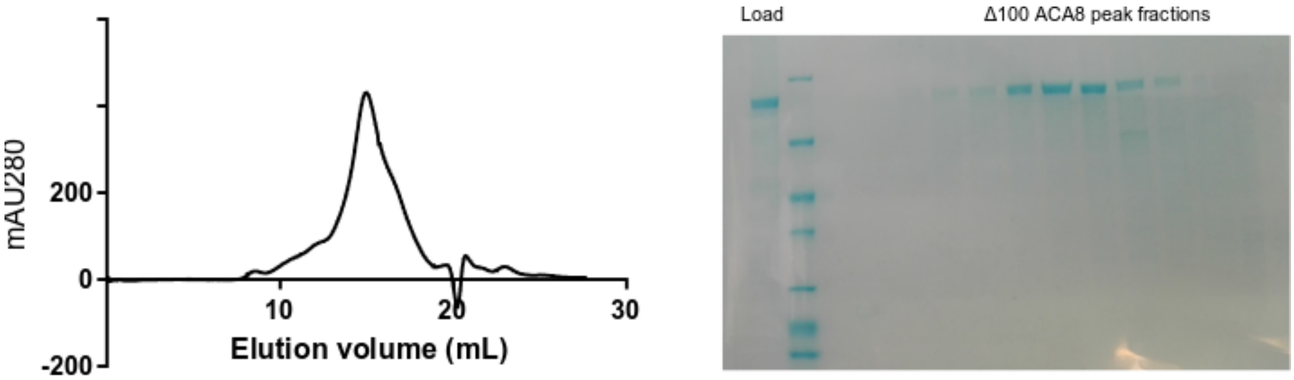

**Figure S7: SEC profile and SDS-PAGE from Δ100-ACA8 purification**

*Left:* The elution profile for Δ100. *Right:* The SDS PAGE of the peak fraction.

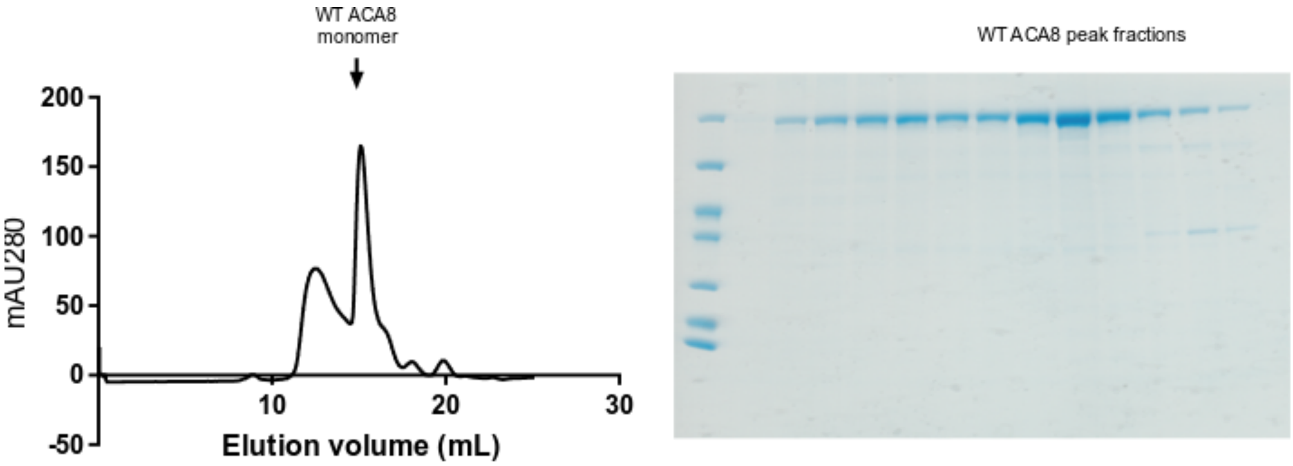

**Figure S8: SEC profile and SDS-PAGE from WT-ACA8 purification.**

*Left:* The elution profile for WT ACA8 with the monomer peak denoted by an arrow. *Right:* The SDS PAGE of the peak fraction.

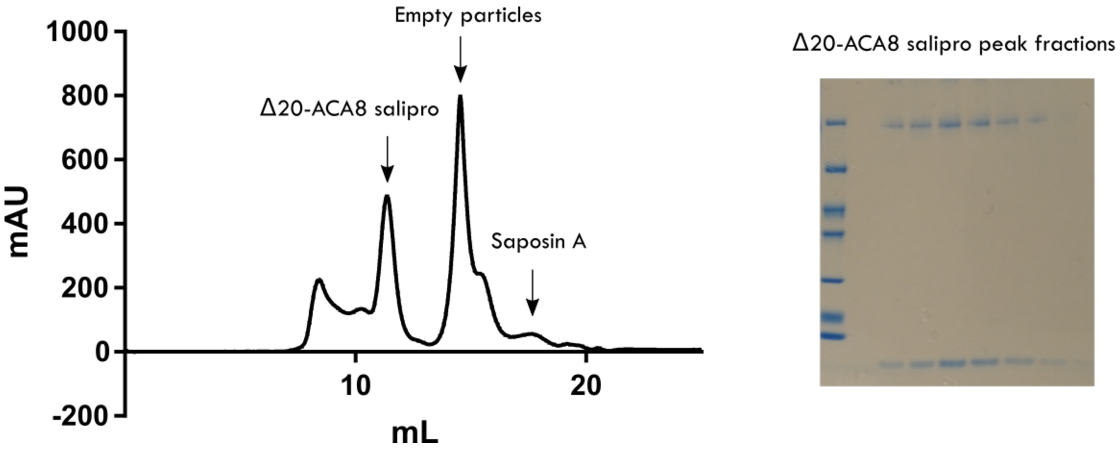

**Figure S9: SEC profile of salipro reconstituted Δ20-ACA8 and a gel representative of peak fractions.** *Left:* The elution profile for salipro reconstituted Δ20-ACA8 with the arrows denoting content of peaks. *Right:* The SDS PAGE of the Δ20-ACA8 salipro peak fraction.

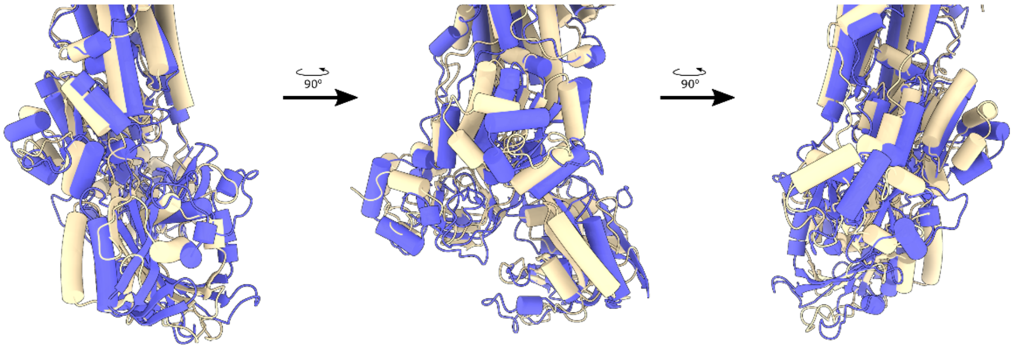

**Figure S10: ACA8 E2P structure compared to SERCA E2P.**

ACA8 is in wheat color, and SERCA (PDB: 3B9B) is in blue. All cytoplasmic domains are vertically shifted to a similar degree.

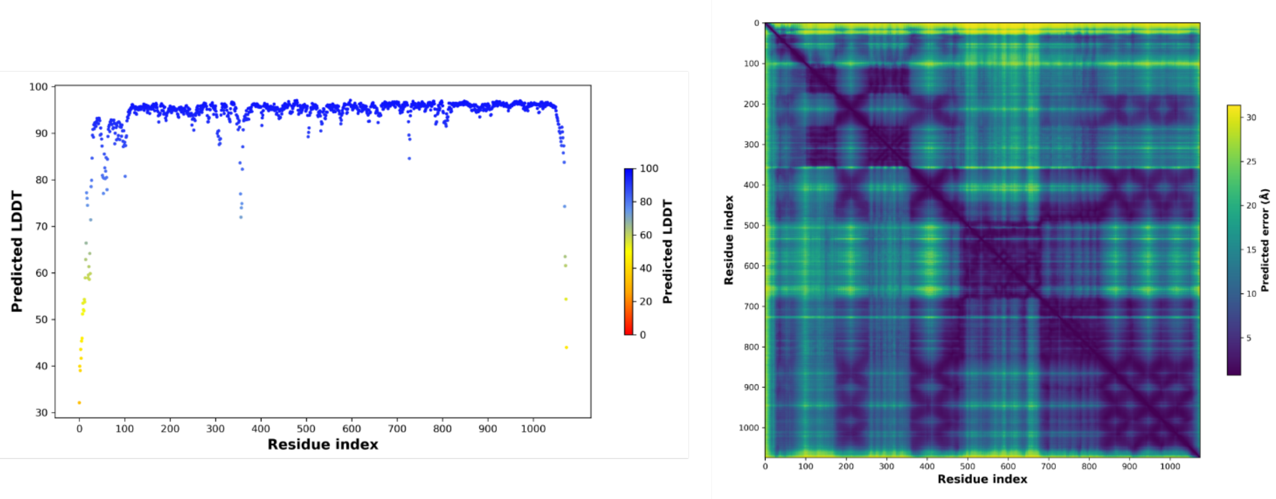

**Figure S11: Confidence level of each residue in the ACA8 AlphaFold prediction.**

*Left:* The pLDDT score for the residues of the AlphaFold predicted ACA8 structure. *Right:* The predicted aligned error for the residues in the AlphaFold predicted ACA8 structure.

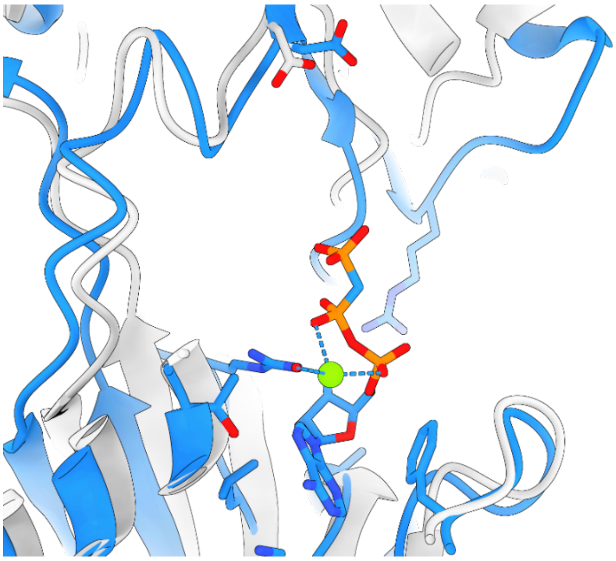

**Figure S12: Modulatory ATP binding can be accommodated in the ACA8 AlphaFold Predicted structure.** ACA8 AlphaFold predicted structure compared to a structure of SERCA with bound modulatory ATP (2C88), aligned on the P domain. The distance to the catalytic Asp is large for modulatory ATP binding.

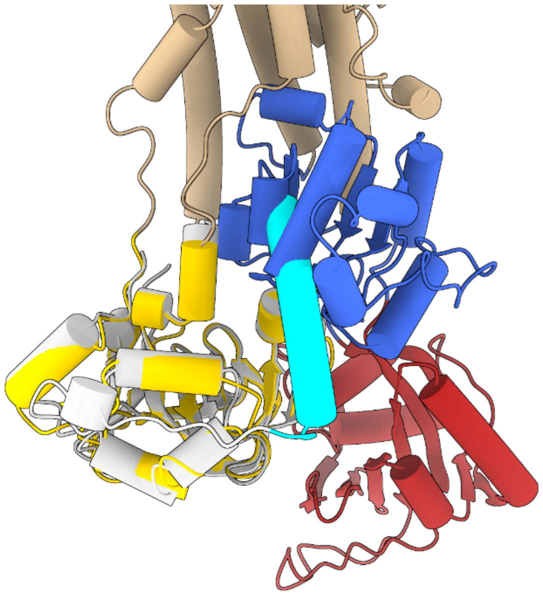

**Figure S13: The extended helix of the autoinhibitory domain hampers the E1 to E2 transition.**

The A domain of the ACA8 AlphaFold predicted model is shown in gray with CaMBS2 of the autoinhibitory domain in cyan aligned with ACA8 E2P state colored by domain, TM domain in wheat, A domain in yellow, P domain in blue and N domain in red. The second CaM binding site of the autoinhibitory domain would clash with the P domain. Transition from E1 to E2P/E2 states is hampered by CaMBS2.

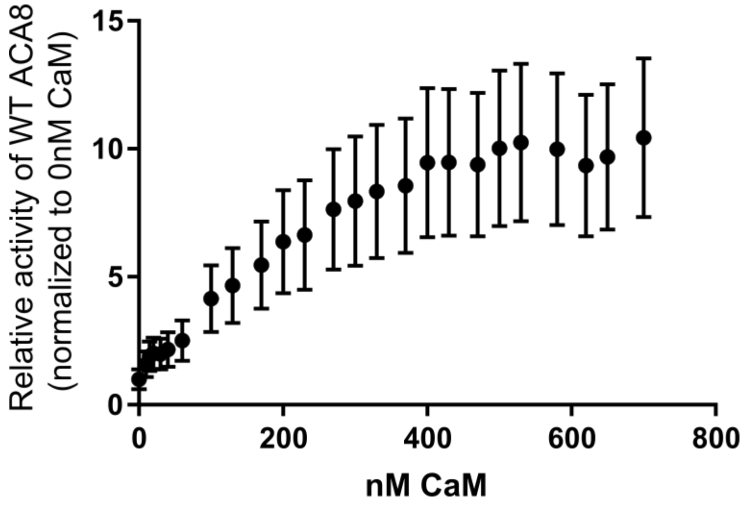

**Figure S14: WT ACA8 activation by calmodulin.**

CaM titration from 0-700nM. The $Ca^{2+}$ ATPase activity was tested as described in the method section. The CaM concentration dependence was tested using 24 different CaM concentration (ranging from 10-700 nM). The activity was tested with and without $Ca^{2+}$. Activities without $Ca^{2+}$ (1 mM EGTA) were subtracted from activities with $Ca^{2+}$. Two experiments in technical triplicates were performed.

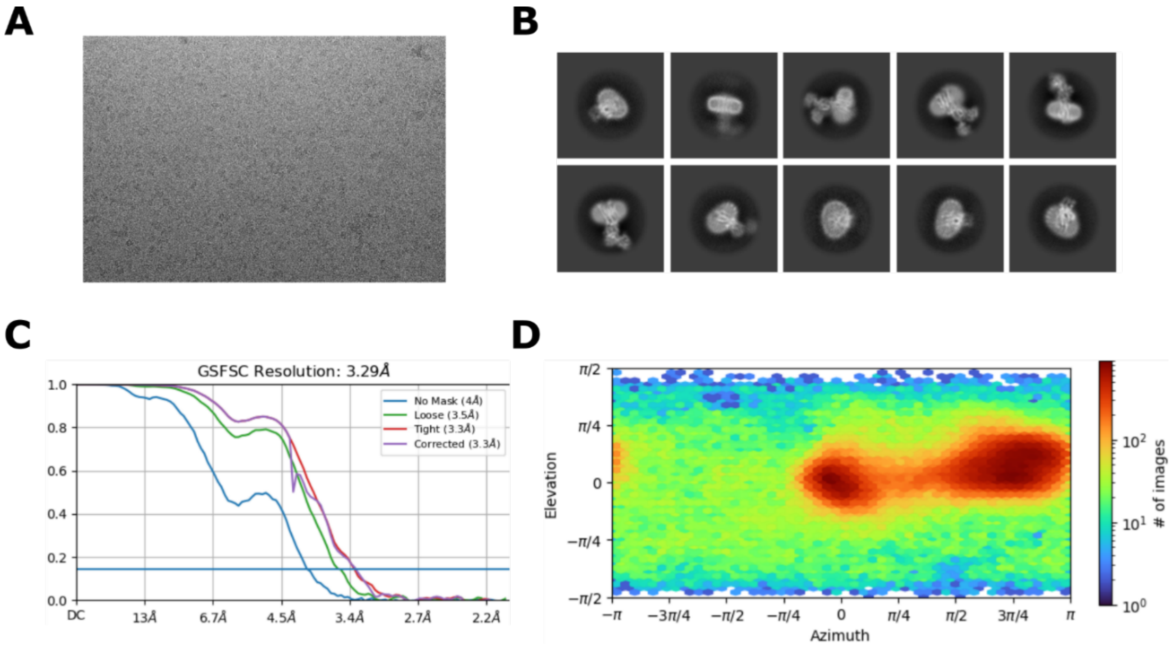

**Figure S15: ACA8 E2P state data processing**

**A)** Representative micrograph from ACA8 E2P dataset. **B)** Representative 2D classifications. **C)** Gold standard Fourier shell correlation curve for the ACA8 E2P data processing. **D)** Viewing direction distribution.

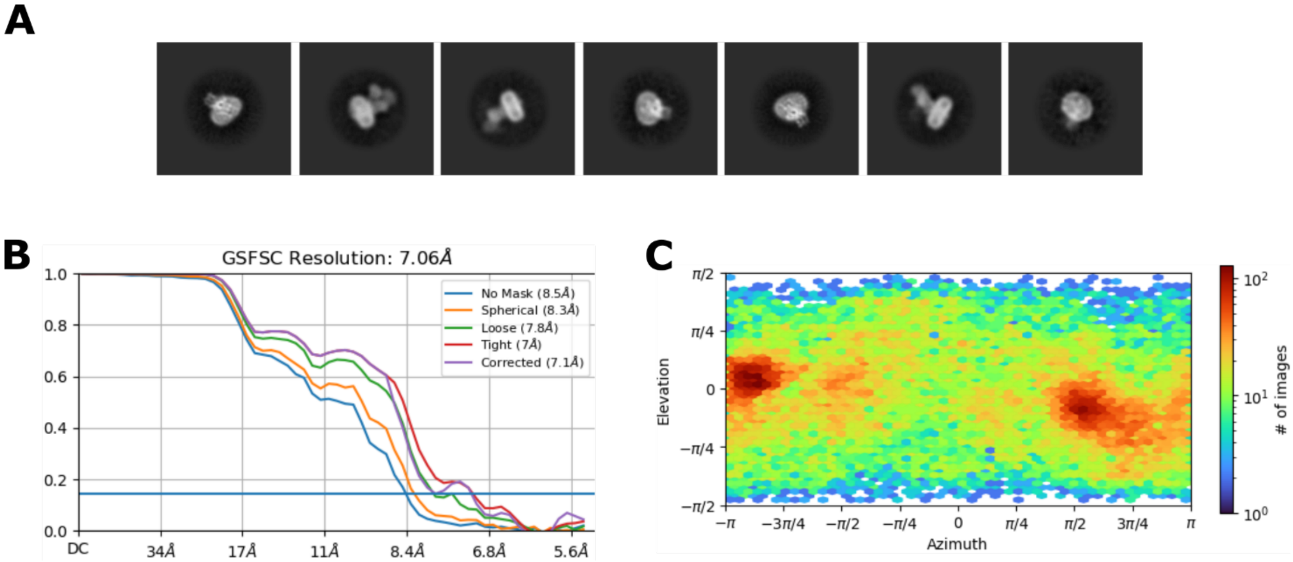

**Figure S16: ACA8 E1 state data processing.**

**A)** Representative 2D classes from the E1 state data. **B)** Gold standard Fourier shell correlation curve for the ACA8 E1 state data processing. **C)** Viewing direction distribution.

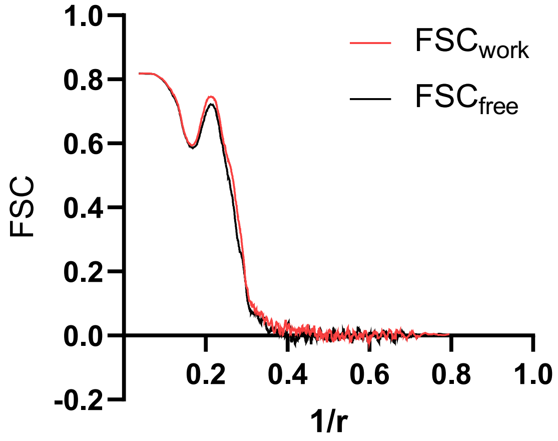

**Figure S17: Cross validation of the ACA8 E2P model.** If FSCwork and FSCfree are very different it indicates overfitting.

**Table S1: Specific activities and fold activation of purified ACA8 ± S.E.** Results are from 6 (WT-ACA8 (8 for $Ca^{2+}$/CaM)), 4 (Δ20-ACA8) and 9 (Δ100-ACA8), 6 (Δ20-ACA8 acidic phospholipids) and 5 (Δ20-ACA8 salipro (9 for $Ca^{2+}$/CaM)) different independent experiments.

| | $Ca^{2+}$ | $Ca^{2+}$/CaM | Fold activation |
|---|---|---|---|
| | *nmol Pi/min/µg ACA8* | *nmol Pi/min/µg ACA8* | *Fold activation* |
| WT-ACA8 | 0.242 ± 0.0446 | 2.251 ± 0.150 | 10.4 ± 1.8 |
| Δ20-ACA8 | 0.0628 ± 0.0024 | 1.193 ± 0.0717 | 19.2 ± 1.8 |
| Δ100-ACA8 | 1.098 ± 0.104 | 1.163 ± 0.125 | 1.06 ± 0.033 |
| Δ20-ACA8 Acidic phospholidpids | 1.55 ± 0.124 | 2.76 ± 0.330 | 1.770 ± 0.130 |
| Δ20-ACA8 salipro | 0.034 ± 0.0089 | 0.864 ± 0.252 | 29.01 ± 3.61 |

**Table S2: Data collection and processing statistics**

| Data collection and processing | E2P state | E1 state<br>Dataset 1 | Dataset 2 |
|---|---|---|---|
| Magnification | ×130,000 | ×130,000 | |
| Voltage (kV) | 300 | 300 | |
| Microscope | Titan Krios G3i (Aarhus University) | Titan Krios G3i (Aarhus University) | |
| Camera | Gatan K3 | Gatan K3 | |
| Physical pixel size (Å/pix) | 0.647 | 0.66 | |
| Electron exposure ($e^{-}$/Å$^2$) | 60.06 | 62.2 | 61.8 |
| Defocus range (µm) | 0.8–1.8 | 0.8–1.8 | |
| Number of movies | 5,297 | | |
| Initial particle images (no.) | 941,952 | 996,849 | 752,205 |
| Final particle images (no.) | 193,419 | 40,686 | |
| Symmetry imposed | C1 | C1 | |
| Map resolution (Å) | 3.29 | 7.1 | |
| FSC threshold | 0.143 | 0.143 | |

**Table S3: CryoEM model refinement and validation statistics.**

| **Refinement and validation** | **E2P state** |
|---|---|
| Model composition | |
| Protein residues | 913 |
| Ligands | $Mg^{2+}$: 1 BeF: 1 |
| *B* factors (Å$^2$, min/max/mean) | |
| c Protein | 1.13/121.61/55.62 |
| Ligand | 34.22/47.49/41.79 |
| R.m.s. deviations | |
| Bond lengths (Å) | 0.003 |
| Bond angles (°) | 0.503 |
| **Validation** | |
| MolProbity score | 1.80 |
| Clashscore | 9.64 |
| Rotamer outliers (%) | 0.00 |
| Ramachandran plot | |
| Favored (%) | 95.82 |
| Allowed (%) | 4.18 |
| Disallowed (%) | 0.0 |
